# Work horse strain *Clostridioides difficile* 630Δ*erm* is oblivious to its anaerobic lifestyle

**DOI:** 10.1101/2020.01.07.897181

**Authors:** Daniel Troitzsch, Hao Zhang, Silvia Dittmann, Dorothee Düsterhöft, Annika-Marisa Michel, Lothar Jänsch, Katharina Riedel, José Manuel Borrero-de Acuña, Dieter Jahn, Susanne Sievers

**Affiliations:** Department of Microbial Physiology and Molecular Biology, Institute of Microbiology, University of Greifswald, Greifswald, Germany; Institute of Microbiology, Technical University of Braunschweig, Braunschweig, Germany; Institute of Biochemistry, University of Leipzig, Leipzig, Germany; Helmholtz Center for Infection Research, Braunschweig, Germany; Braunschweig Integrated Centre of Systems Biology (BRICS), Technical University of Braunschweig, Braunschweig, Germany

## Abstract

The laboratory reference strain 630Δ*erm* of the anaerobic human pathogen *Clostridioides difficile* is characterized by a remarkable high oxygen tolerance. We show that an amino acid exchange in the DNA binding domain of the hydrogen peroxide sensor PerR results in a constitutive derepression of PerR-controlled genes and thus in an oxidative stress response even under anaerobic conditions. This questions the model status, strain 630Δ*erm* claims in *C. difficile* research.

## Introduction

*Clostridioides difficile* (*C. difficile*) is a Gram-positive, anaerobic, spore-forming pathogen causing primarily hospital-acquired, but increasingly also community-acquired infections, which turned the bacterium into one of the most problematic pathogens in human health care nowadays. *C. difficile* infections (CDIs) are often associated with broad-spectrum antibiotic therapy. Clinical symptoms of CDI vary from light diarrhea to acute infections like pseudomembranous colitis (1).

Due to its anaerobic lifestyle, oxygen (O_2_) and reactive O_2_ species in the human intestine represent a challenge for *C. difficile*. Remarkably, a high tolerance to O_2_ was recently reported for a sporulation deficient mutant of *C. difficile* 630Δ*erm* (2). Strain 630Δ*erm* is an erythromycin-sensitive and laboratory-generated derivative of the original patient-isolated strain 630 and is commonly used by *C. difficile* researchers as reference strain for the generation of gene knock out mutants (3). Although the oxidative stress response is vital for an intestinal pathogen infecting its host, knowledge on the molecular details of oxidative adaptation mechanisms in *C. difficile* is still limited (4, 5).

## Results and Discussion

In previous studies, we observed a high abundance of oxidative stress-related proteins in *C. difficile* 630Δ*erm* already at conditions devoid of any oxidizing agents (6) and no significant induction when the bacterium was shifted to micro-aerobic conditions (7). Several of the corresponding genes are encoded at one genetic locus comprising a rubrerythrin (*rbr1*), the transcriptional repressor PerR (*perR*), a desulfoferrodoxin (*rbo*) and a glutamate dehydrogenase with an N-terminal rubredoxin fold (*CD630_08280*). Rbr1 even represents the second most abundant protein after the S-layer protein SlpA (6). We enquired, why these genes are highly expressed in the absence of any oxidative stress and focused on the repressor protein PerR, which regulates its own transcription and the one of genes involved in oxidative stress and metal homeostasis as described in *Bacillus subtilis* and in other Gram-positive bacteria (8, 9). PerR is a member of the ferric uptake regulator (Fur) family and senses H_2_O_2_ stress by metal-catalyzed histidine oxidation (10), (Fig. 1A). Due to the permanently high cellular concentration of proteins encoded in the *rbr1* operon, we hypothesized a constitutive expression of the operon possibly caused by failure of PerR-mediated gene repression under anaerobic conditions.

**Figure 1:**
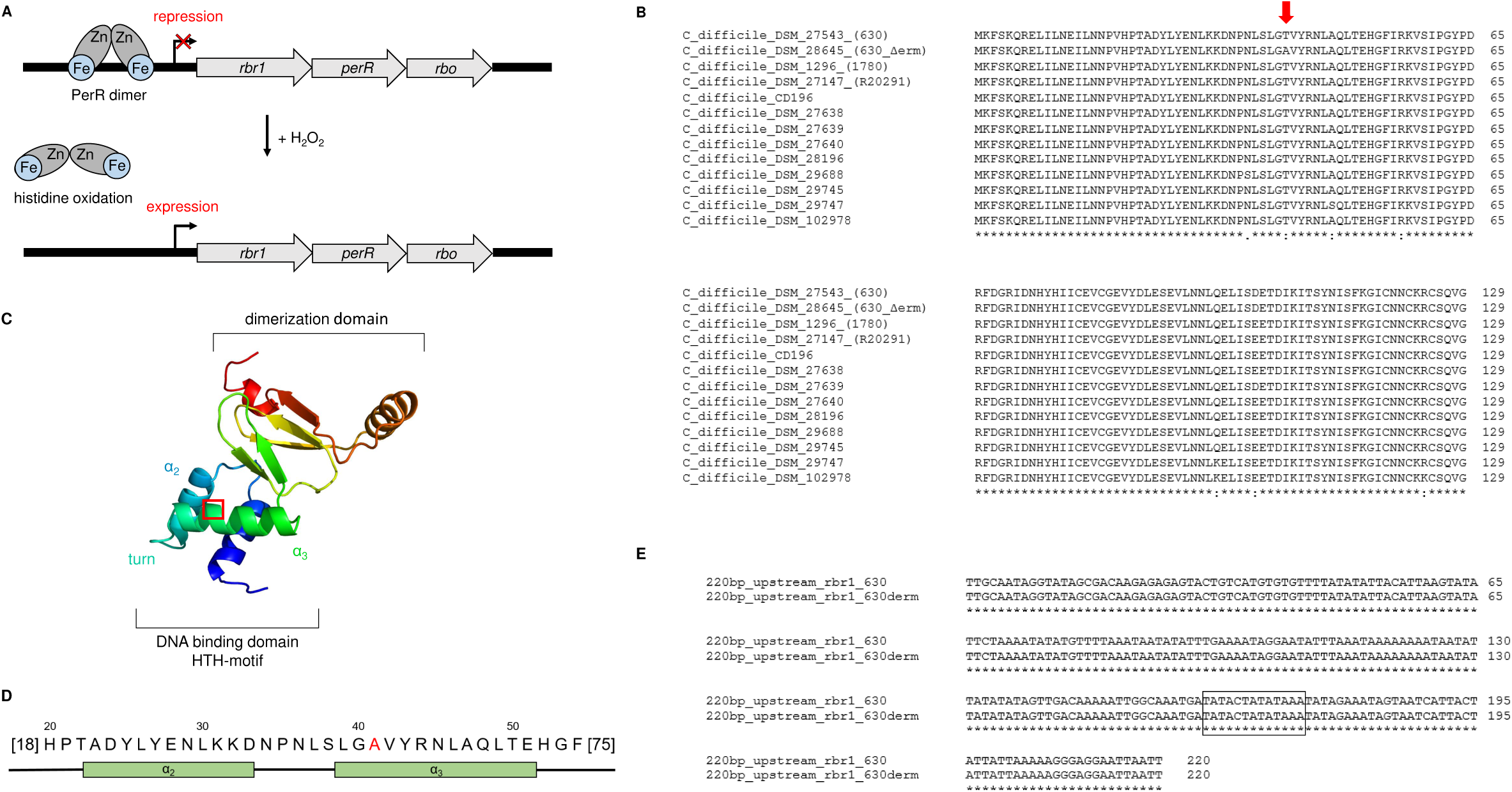
DNA binding of PerR. A] Mode of PerR function is shown schematically. PerR bound to the DNA represses expression of the *rbr1* operon. H2O2 treatment leads to PerR oxidation, a conformational change and release of the DNA promoter resulting in *rbr1* expression. B] Alignment of amino acid sequences of the PerR protein of 13 different *C. difficile* strains. C] Structure of *C. difficile* 630Δ*erm* PerR deduced from the homolog of *B. subtilis*. The DNA binding site including the HTH-motif and the dimerization domain are marked. D] Amino acid sequence of the DNA binding site of *C. difficile* 630Δ*erm* PerR. The T41A mutation is marked red. E] Alignment of the promoter sequences of the *rbr1* gene in *C. difficile* strains 630 and 630Δ*erm*.

It was reported before, that the lab-generated strain *C. difficile* 630Δ*erm* features several genome alterations compared to its parental strain *C. difficile* 630 (11, 12). We aligned *perR*-sequences of *C. difficile* 630 and *C. difficile* 630Δ*erm* and found a single nucleotide polymorphism (SNP, A > G), resulting in an amino acid conversion from threonine to alanine at position 41. An alignment of the PerR amino acid sequence with 11 other clinically relevant *C. difficile* strains revealed that the T41A substitution is unique to the laboratory strain 630Δ*erm* (Fig. 1B). A comprehensive sequence alignment of over 900 proteins of the Fur family from different species showed that the threonine in position 41 is highly conserved and present in over 80% of the investigated proteins. More than 90% of the proteins contain a threonine or serine at this position (Dataset S1). A structural comparison of previously investigated DNA binding domains of Fur and PerR homologues in *Escherichia coli, B. subtilis, Staphylococcus epidermidis* and *Streptococcus pyogenes* to the *C. difficile* 630Δ*erm* PerR sequence indicates the T41A mutation to be located in a helix of the helix-turn-helix motif of the DNA binding domain (Fig. 1C and D). The DNA promoter sequences upstream of *rbr1* are identical between strain 630 and 630Δ*erm* (Fig. 1E). We therefore hypothesized that the amino acid substitution in PerR is the reason for loss of binding of the repressor to PerR boxes on the DNA and possibly causes increased O_2_ tolerance of strain 630Δ*erm* compared to other *C. difficile* strains including its parental strain 630. To investigate differences in O_2_ tolerance between *C. difficile* 630 and 630Δ*erm* we counted colony forming units (CFU) for both strains after cells have been exposed to atmospheric O_2_ concentrations (Fig. 2A). Strain 630 showed a significantly higher susceptibility to O_2_ than its derivative 630Δ*erm*, of which a substantial number of cells survived even after 9 h of challenge.

**Figure 2:**
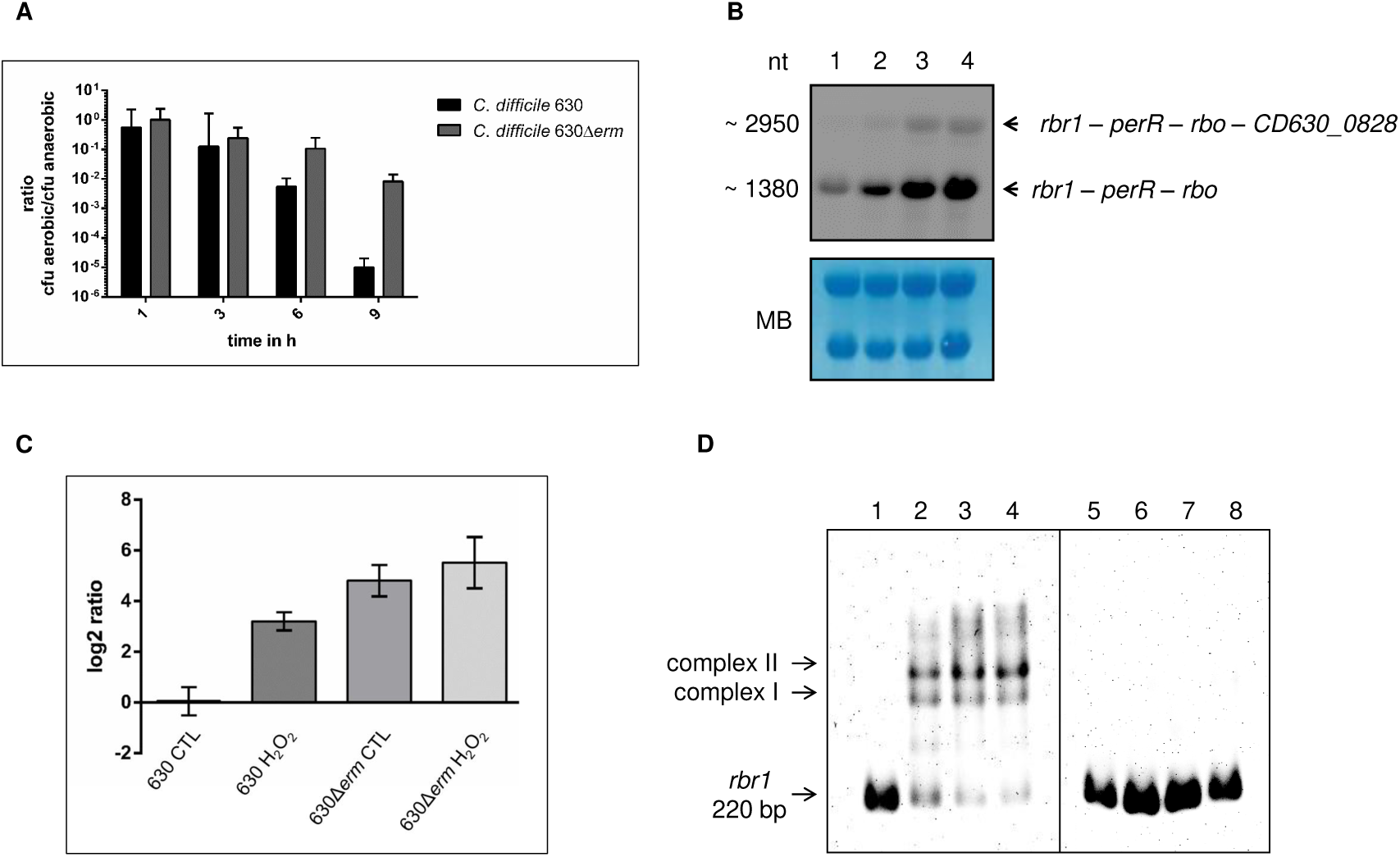
CFU counting, transcriptional analyses and DNA-PerR interaction. A] Survival of *C. difficile* strains 630 and 630Δ*erm* in the presence of O_2_. Cell numbers are related to the cell number determined before the aerobic shift. B] Northern blot analysis of expression of *rbr1* operon: 630 control (1) and 630 H_2_O_2_ induced (2), 630Δ*erm* control (3) and 630Δ*erm* H_2_O_2_ induced (4). Total RNA levels were monitored by methylene blue staining (MB). C] Transcription of the *rbr1* gene was quantified by RT-qPCR analysis and related to *C. difficile* 630 control. D] EMSA analyses were carried out with 2 ng of the *rbr1* promoter fragment (220 bp), concentrated 147 nM, from strain *C. difficile* 630. In lane 1 and 5, the DNA fragment was incubated without protein. The DNA fragment was incubated with purified PerR 630 (lane 2 to 4) or PerR 630Δ*erm* (lane 6 to 8) with increasing protein amount (360 ng, 420 ng and 480 ng, respectively).

To prove that the higher O_2_ tolerance of *C. difficile* 630Δ*erm* was caused by the missing binding of PerR to its cognate DNA binding site, we analyzed transcriptional levels of the *rbr1* operon under hydrogen peroxide (H_2_O_2_) stress and control conditions (Fig. 2B and C). Transcription of the operon was inducible by H_2_O_2_ in strain 630 by a factor of 9, whereas transcript levels in 630Δ*erm* were permanently very high and almost not inducible by H_2_O_2_ (factor of 1.6).

To confirm that the T41A exchange in PerR of strain 630Δ*erm* is the sole reason that hampers PerR-box binding, we performed electrophoretic mobility shift assays (EMSA) excluding any other cellular factors. PerR proteins from strains 630 and 630Δ*erm* were recombinantly produced, purified and incubated with a labelled 220 bp upstream promoter fragment of *rbr1*. While PerR from strain 630 led to a clear shift of the DNA band, no shift was detectable for PerR from *C. difficile* 630Δ*erm* at any tested protein concentration (Fig. 2D).

This study demonstrated the constitutive derepression of genes involved in the oxidative stress response in strain *C. difficile* 630Δ*erm* caused by only one SNP in the DNA sequence of the transcriptional repressor PerR. Since this strain is used as reference in many laboratories for the construction of gene inactivation mutants, researchers should be aware of its permanent oxidative stress response.

## Materials and Methods

### Bioinformatic methods

Structural analysis of the *C. difficile* PerR protein was performed using the Phyre2 web portal (13). For the alignment of over 900 Fur family proteins MView was used (14). The alignment of promoter sequences was carried out with Clustal Omega (15). PerR Boxes were identified using Virtual Footprint Version 3.0 (16).

### Bacterial strains and growth conditions

*C. difficile* strains were obtained from the German Collection of Microorganisms and Cell Cultures GmbH (DSMZ, Braunschweig, Germany) and cultured as previously described (17). For Northern Blot and RT-qPCR analyses, *C. difficile* 630 and *C. difficile* 630Δ*erm* were grown to an A_600_ of 0.4, before cultures were split and one of the two subcultures stressed with 0.4 mM H_2_O_2_ for 10 min. Samples were taken to allow for a later preparation of RNA (18). For CFU counting experiments, 10 mL of the *C. difficile* culture at A_600_ of 0.4 were transferred to a 92 × 16 mm petri dish and aerobically incubated. Samples were taken before and 1, 3, 6 and 9 h after oxygen exposure in three biological replicates, and dilution series incubated anaerobically. For overexpression of PerR, *Escherichia coli* BL21 grown in LB medium was used.

### RNA preparation

For cell lyses and RNA isolation TRIzolTM Reagent provided by Invitrogen (Thermo Fisher Scientific; Waltham, Massachusetts, USA) was used according to the manufacturer’s protocol (19). RNA solubilized in DEPC-treated water was stored at −70 °C.

### Transcriptional profiling

A PCR fragment of gene *rbr1* was prepared using chromosomal DNA of *C. difficile* 630 as a template with primers 5’-AATGGCAGGATTTGCAGGAG-3’ and 5’-CTAATACGACTCACTATAGGGAGATGGATGGTCACATACTGGGC-3’. Digoxygenin (DIG)-(18) labeled RNA probes were obtained and Northern Blot analyses carried out as previously described (18). *Rbr1* transcription was quantified by RT-qPCR in three biological replicates with three technical replicates each using above mentioned primers. The *codY* gene with forward primer 5’-ATTAGGAACATTGGTACTTTCAAGAT-3’ and reverse primer 5’-TTGAACTACAGCTTTCTTTCTCATT-3’ served as reference. cDNA synthesis and qPCR was performed as described elsewhere (20). The qPCR reactions were performed on a qTOWER 2.2 quantitative PCR thermocycler (Analytik Jena). Quantitative data analysis was based on the Pfaffl method (21).

### Overexpression and purification of PerR from *C. difficile* 630 and *C. difficile* 630Δ*erm*

Slightly modified guidelines for protein overproduction were followed as previously reported (22). Firstly, synthesized gene variants (ThermoFisher, Darmstadt, Germany) were introduced into the pGEX6P1 allowing for GST-mediated affinity purification and subsequent tag excision. *E. coli* BL21 cells harboring recombinant genes were induced (0.1 mM IPTG) at an OD_600nm_ of 0.5, grown aerobically for 4 h and shifted to anaerobiosis for 2h.

### Electrophoretic mobility shift assay (EMSA)

Shift assays were conducted as previously specified (23) with minor variations. The 220 bp *rbr1* promoter region was amplified via PCR using forward primer 5’-TTGCAATAGGTATAGCGACAAG-3’ and reverse primer 5’-TGCAATAGGTATAGCGACAAG-3’. The EMSA was performed under anaerobic conditions.

## Acknowledgments

This work was funded by the German Research Foundation (231396381/GRK1947), the „Ministerium für Bildung, Wissenschaft und Kultur Mecklenburg-Vorpommern” (UG16001), the Federal State of Lower Saxony, Niedersächsisches Vorab CDiff and CDInfect projects (VWZN2889/3215/3266).

## Legend of Dataset S1

### Dataset S1

**Sequence alignment of Fur family proteins of different bacteria**

Sequences of over 900 proteins of the Fur family were aligned using a PSI-Blast via Phyre2 (13). Conserved amino acid residues are indicated at the end of the table.

