## Supplementary figures and images for "Work horse strain *Clostridioides difficile* 630Δ*erm* is oblivious to its anaerobic lifestyle"

### Supplemental Dataset 1

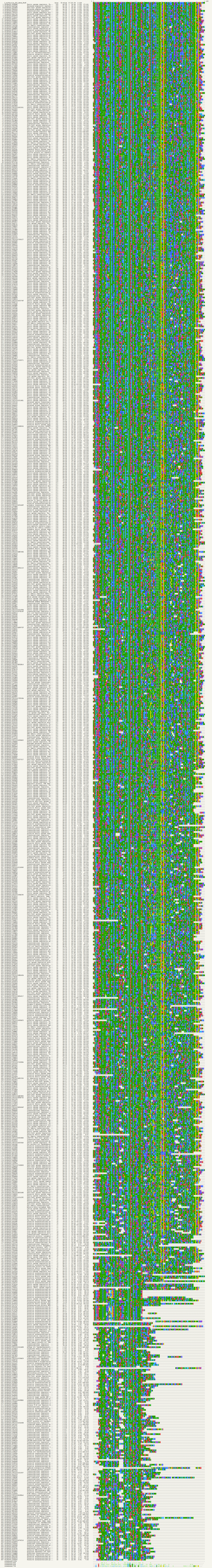
